# Dorsomedial prefrontal cortex acts as an integrative hub during information gathering

**DOI:** 10.64898/2026.08.11.744102

**Authors:** K. Kadri, A.A. Marzuki, M. del Rio, T.U. Hauser

**Affiliations:** Department of Psychiatry and Psychotherapy, Faculty of Medicine, University Tuebingen, Tübingen, Germany; Functional Imaging Laboratory (FIL), Department of Imaging Neuroscience, University College London, London, UK; Max Planck UCL Centre for Computational Psychiatry and Ageing Research, University College London, UK; German Center for Mental Health (DZPG), Partner Site Tübingen, Tübingen, Germany

## Abstract

Gathering information before committing to a choice is critical in real-world decision making, and biases thereof are hallmark features of psychiatric disorders. Here, we study the behavioural and neural mechanisms that guide information gathering and characterise several key cognitive constituents, including escalating urgency and systematically biased temporal weighting of information. Using fMRI, we identify an integrated information gathering signal in ventromedial and dorsomedial prefrontal cortices (dmPFC), signalling an overall likelihood for continued sampling of information. Teasing this signal apart, we find distinct neural circuits encoding separable information-gathering constituents: whilst an urgency signal primarily engaged locus coeruleus and dmPFC, accumulated evidence was represented in anterio-medial PFC, and evidence-strength prediction errors were computed in ventral striatum and dmPFC. These findings indicate that information gathering arises from functionally distinguishable prefrontal-subcortical computations that converge within medial prefrontal cortex, providing a mechanistic framework for understanding aberrant sampling in psychiatric conditions, including schizophrenia and obsessive-compulsive disorder.

## Introduction

Many decisions permit a delay before commitment, allowing additional information to be gathered about candidate options. Such sampling can improve choice but incurs time, effort, and opportunity costs. Deciding when to cease gathering information and commit to a choice therefore constitutes a non-trivial arbitration process ^1,2^.

Information gathering has traditionally been examined using behavioural paradigms, including beads and urns tasks ^3^, which reveal reliable and marked individual differences ^4,5^, often linked to mental health ^6–11^. Much less is known about how the brain solves the conundrum of when to decide. Prior non-mechanistic research has implicated the dorsal prefrontal cortex ^12,13^, particularly structures along the dorsomedial wall, including anterior cingulate cortex (ACC; Furl C Averbeck, 2011; Kobayashi & Kable, 2024).

More recent work using advanced behavioural paradigms and modelling has revealed that information gathering is a complex cognitive process where distinct processes are arbitrated ^4,9^, potentially subserved by diverse neurotransmitter systems ^5,16^. Firstly, gathered information seems to be accumulated non-linearly, with a pronounced recency bias ^4^. Participants consistently overweight recent evidence while underweighting equally relevant earlier information. This suboptimal weighting resembles biases in perceptual decision making ^17,18^ and is consistent with a prediction-error-like learning process^19^, whereby incoming information is evaluated against expectations and an evidence-strength prediction error drives sampling cessation^4^.

Secondly, information gathering is also guided by information-independent factors, including a gradually emerging urgency signal whereby individuals become increasingly willing to commit as time elapses, irrespective of evidence ^4,9,20^. This process is akin to urgency signals in perceptual decision making ^21,22^ and collapsing boundaries in sequential-sampling models ^20,23^. These urgency-like signals in perceptual decision making are thought to be encoded in subcortical-prefrontal networks ^24–27^, but have not been examined during information gathering, where such analogue signals emerge over substantially slower timescales.

To determine whether distinct cognitive constituents of information gathering show separable neural underpinnings and how they are integrated into a common decision-commitment signal, we developed a paradigm that disentangles mechanistic contributors to information gathering. Using fMRI, we show that these constituents recruit distinct subcortical-prefrontal circuits but eventually converge in dorsomedial prefrontal cortex (dmPFC) to form a unified decision signal. Our findings thus demonstrate how integrating across multiple decision-relevant neural processes culminates in unified information-gathering behaviour, critical for understanding biases associated with psychiatric conditions.

## Results

To assess neural mechanisms driving information gathering, we recruited 29 healthy participants (13 females; mean age ± s.d., 27.83 ± 7.99 years) who completed an fMRI version of an information-gathering task (160 trials; Fig. 1A) adapted from a previously developed paradigm ^4^.

**Figure 1:**
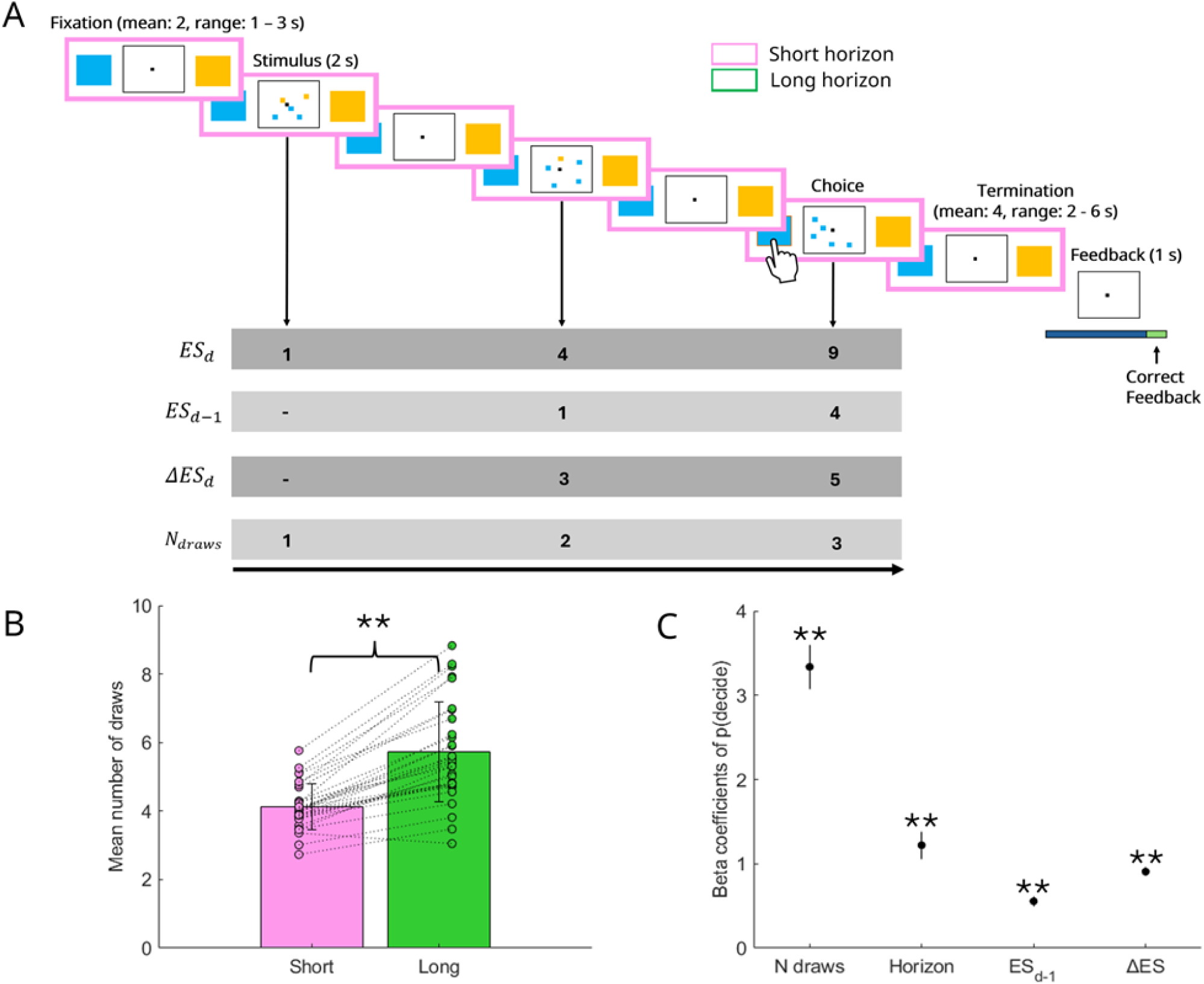
Separable cognitive contributors influence information gathering. A. Participants completed an fMRI information-gathering task in which they decided which stimulus (yellow or blue gems) was more abundant on each game. A sequence of draws was presented until participants committed to a decision; each draw comprised five stimuli in varying proportions. Evidence strength (ES) at draw d was quantified as the cumulative difference in evidence favouring the gem that was more abundant at draw d. Cumulative evidence strength at the previous draw (ESd-1) was defined as ES lagged by one draw, and an evidence-strength prediction error (ΔES) as the signed difference between cumulative ES at draw d-1 and draw d. Horizon length was cued by frame colour before stimulus display. B. Participants gathered more information during the long horizon. Error bars indicate s.e.m. C. A generalised linear mixed model showed that urgency (number of draws), shorter horizons, ESd-1, and ΔES significantly predicted cessation of information gathering and decision commitment [p(decide)]. Error bars indicate s.e.m. **p < .01.

On each trial, participants decided which of two stimuli (yellow or blue) was more abundant in a large arena. They observed a continuous sequence of draws (five items per draw) until committing to a choice. Two-time horizons constrained sampling: a short horizon (4-8 samples) and a long horizon (8-12 samples). Participants did not know the exact sampling limit within each horizon, and if they had not made a decision by the predetermined endpoint, the trial terminated and they incurred a small loss.

### Separable cognitive mechanisms determine information gathering

To establish task engagement, we first examined behavioural performance. Participants selected the more abundant colour in 73.97% ±2.92) of trials, incorrect colours in 21.12% (± 3.81), and timed out in 4.91% ± 4.30, indicating that displayed evidence guided choices. They were also sensitive to horizon length, gathering less information in short than long horizons (short, 4.12 ± 0.66 draws; long, 5.72 ± 1.46 draws; t(29) = -9.38, 95% CI [-1.95, -1.25], p < .001; Fig. 1B).

To formally quantify the constituents of information gathering, we used model developed for this task class ^4,8^, which captured how experimental and information-related factors determined whether participants committed to a choice or continued sampling (p(decide); Methods).

The horizon condition significantly affected choice commitment, mirroring above behavioural finding (b = 1.22, SE = 0.165, p < .001; Fig. 1C). The number of draws, indexing an urgency signal as sampling progressed ^4,8,9^, also exerted a strong positive effect (b = 3.34, SE = 0.259, p < .001; Fig. 1C), indicating that participants became increasingly willing to decide as they sampled more information, independently of evidence.

To assess how information itself guided commitment, we operationalised evidence by separating accumulated prior evidence (evidence favouring the majority at the previous sample: ESd-1) from an evidence-strength prediction error (ΔES), reflecting the deviation of current information from ESd-1 ^4^. Both ESd-1 (b = 0.556, SE = 0.061, p < .001) and ΔES (b = 0.908, SE = 0.052, p < .001) positively predicted p(decide). Critically, ΔES exerted a stronger influence than ESd-1 (b = 0.908 vs 0.556), showing that participants over-weighted most recent evidence and capitalised on a prediction-error-like information accumulation process.

### dmPFC and locus coeruleus signal cessation of information gathering

Having established the behavioural constituents of information gathering, we next assessed neural circuitry underlying an integrative signal that drives sampling cessation. We extracted trial-wise predictions indexing participants’ subjective tendency to commit, termed the decision-commitment signal (DCS). DCS was represented in two hypothesised regions: dmPFC (right, MNI = [6 39 22], t = 12.67; left, MNI = [-4 33 20], t=10.16, cluster-extent FWE-corrected p < .05, extending to the ACC; Fig. 2A; Table 1) and bilateral locus coeruleus (LC; right, MNI = [6 -32 -18], t = 5.85, FWE p < .001; left, MNI = [- 6 -30 -14], t = 4.47, pFWE < .001; confirmed via small volume correction (SVC) using an anatomical LC mask, cluster-level FWE-corrected p < .05; see Method). Activity scaled linearly with DCS in dmPFC (Fig. 2B; b = 2.07, 95% CI [1.81, 2.33], p < .001) and bilateral LC (b = 0.31, 95% CI [0.07 0.55], p = .012), and remained significant after controlling for potential confounds, such as button presses or horizon length (Supplementary Figs. S1-S3).

**Figure 2:**
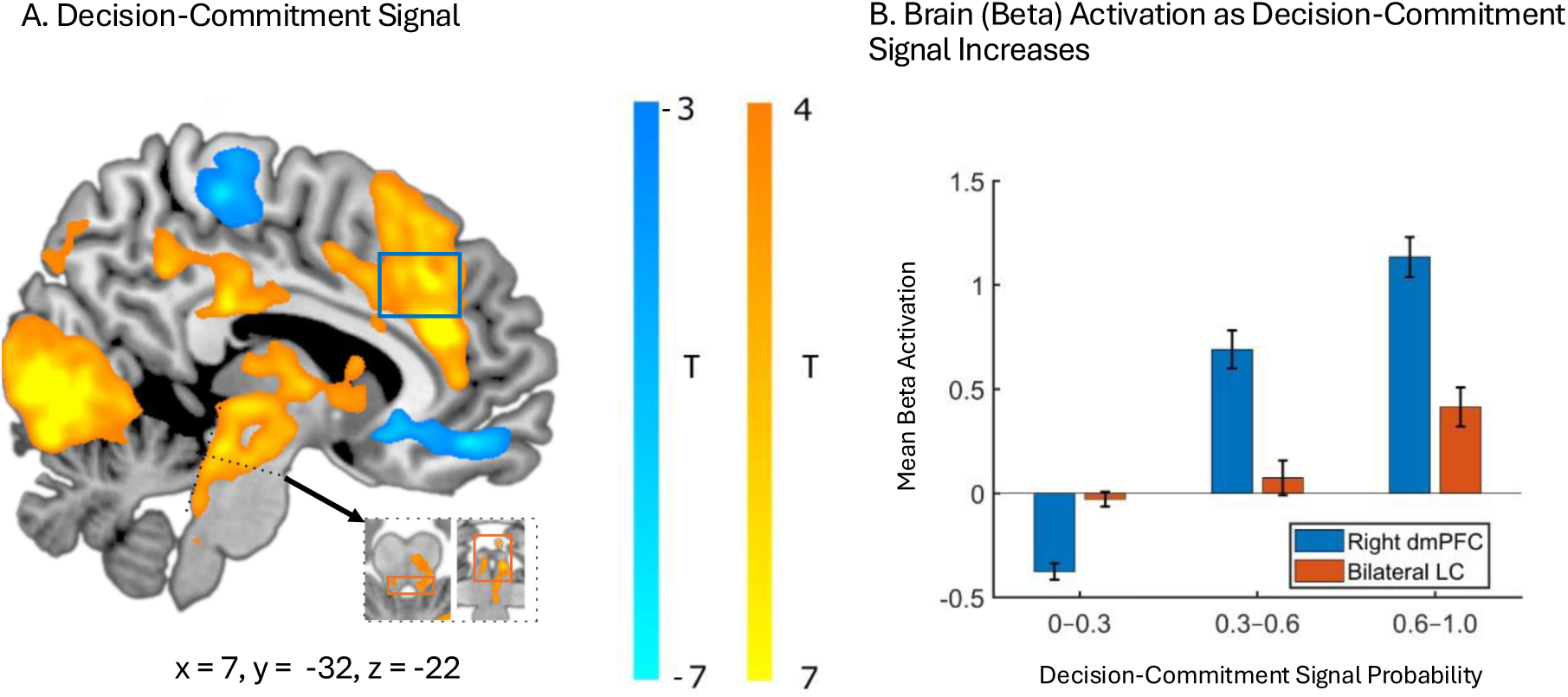
Neural correlates of overall information gathering: decision-commitment signal. A. Decision-commitment (yellow) was encoded in right dmPFC (MNI = [c 3S 22]) and bilateral LC (right, MNI = [c -32 -18]; left, MNI = [-4 -37 -22]), both regions showing cluster-extent FWE-corrected p < .001. Lowered DCS (blue blobs) was associated with activation in right premotor area (MNI = [c -28 c3]) and right vmPFC (MNI = [8 45 -12]). B. Activation in right dmPFC and bilateral LC both showed a linear increase as the decision-commitment signal increased (extracted from a 3mm-radius sphere centred on MNI = [c 3S 22], for illustrative purposes only). Blue and orange squares in 2B correspond to right dmPFC and bilateral LC regions, respectively. Normalised mean beta activity values are presented on the y-axis.

**Table 1.** Peak activation coordinates. Clusters were defined using a voxel-wise threshold of p < .001 (uncorrected), with cluster-level FWE correction at p < .05 unless otherwise indicated. * pFWE (peak) < .05; pFWE is the family-wise error corrected p-value for a single peak voxel ** SVC pFWE (peak) < .05 (LC: Mäki-Marttunen C Espeseth, 2020; NAcc: Pauli et al., 2018); ‡ p < .0001 (uncorrected), Abbreviations: dmPFC = dorsomedial prefrontal cortex; vmPFC = ventromedial prefrontal cortex; LC = locus coeruleus; aPFC = anterior prefrontal cortex; NAcc = Nucleus Accubens

| Contrast | Region | Hemisphere | Cluster Size (voxels) | X | Y | Z | T Score |
| --- | --- | --- | --- | --- | --- | --- | --- |
| **DCS** | dmPFC* | Right | 1422 | 6 | 39 | 22 | 12.67 |
|  |  | Left |  | -4 | 33 | 20 | 10.16 |
|  | LC** | Right | 2 | 6 | -38 | -24 | 3.66 |
|  |  |  | 1 | 4 | -34 | -16 | 3.50 |
|  |  |  | 1 | 4 | -36 | -22 | 3.45 |
| **-DCS** | vmPFC | Right | 3411 | 8 | 45 | -12 | 6.58 |
|  |  | Left |  | -4 | 44 | -14 | 7.17 |
|  | Pre-motor cortex | Right | 1979 | 6 | -28 | 63 | 7.24 |
| **ES<sub>d-1</sub>** | dmPFC | Right | 9005 | 15 | 60 | 34 | 5.78 |
|  | aPFC | Left |  | -14 | 48 | 42 | 6.44 |
|  | vmPFC | Right | 212 | 8 | 57 | -21 | 3.72 |
| **ΔES** | dmPFC | Right* | 82 | 10 | 44 | 50 | 9.03 |
|  |  | Left* | 30 | -9 | 50 | 26 | 7.70 |
|  | vmPFC | Right‡ | 9657 | 3 | 44 | -26 | 5.58 |
|  |  | Left‡ | 127 | -6 | 60 | -15 | 5.42 |
|  | NAcc | Right** | 104 | 12 | 12 | -9 | 5.49 |
|  |  | Left** | 124 | -12 | 4 | -14 | 7.31 |
| **-ΔES** | dmPFC/preSMA | Left | 957 | -27 | 2 | 54 | 7.33 |
| **Urgency** | dmPFC | Right* | 107 | 8 | 38 | 22 | 11.23 |
|  | LC | Right** | 2 | 4 | -34 | -20 | 4.62 |
|  |  |  | 4 | 6 | -36 | -26 | 4.24 |
|  |  | Left** | 7 | -4 | -36 | -21 | 5.23 |

To identify neural signals supporting continued sampling, we examined the inverse of DCS, which corresponded to the probability of not committing to a choice and continuing the sampling (1-p(decide)). This continuation signal was represented primarily in vmPFC (right, MNI = [8 45 -12], t = 6. 58; left, MNI = [-4 44 -14], t = 7.17; pFWE < .001; Fig. 2A) and right premotor cortex (MNI = [6 -28 63], t = 7.24, pFWE < .001), indicating that these regions track sustained information gathering.

### A designated fronto-striatal network carries evidence-strength prediction errors

To determine whether cognitive constituents of information gathering rely on shared or distinct circuitry, we entered them (urgency, ESd-1, and ΔES) into a single fMRI model, allowing them to compete for explained variance (see Methods). To probe neural representations of gathered information, we separated accumulated prior evidence (ESd-1) from the evidence-strength prediction error (ΔES), which indexes surprise elicited by currently observed information.

Firstly, accumulated prior evidence (ESd-1) elicited widespread activity along the medial prefrontal wall, centred on left anterior prefrontal cortex (aPFC; MNI = [-14 48 42], t = 6.44, pFWE < .001; Fig. 3A, extending from vmPFC (MNI = [8 57 -21], t = 3.72, p < .001) to dmPFC (MNI = [15 60 34], t = 5.78, FWE p < .001; Fig. 3A). dmPFC activity scaled linearly with accumulated prior evidence (b = 0.18, 95% CI [0.12, 0.24], p < .001; Fig. 3B). These regions are well known for valuating the reward of choice options ^28,29^, which we here extend to the context of information valuation.

**Figure 3:**
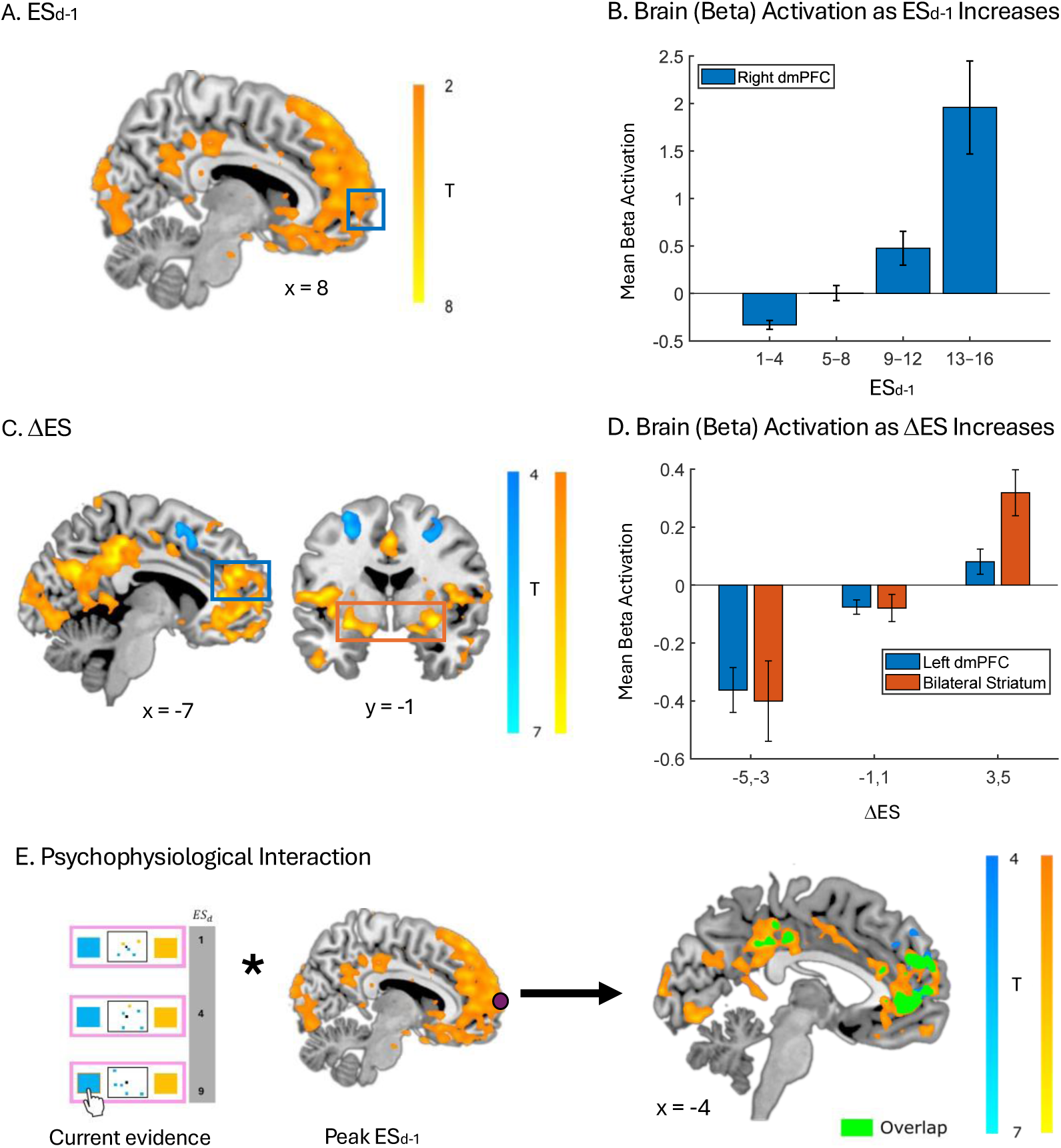
Neural correlates and connectivity underlying accumulated prior evidence (ESd-1) and evidence-strength prediction errors (ΔES). A. ESd-1 was associated with increased dmPFC activation (MNI = [15 c0 34]) and B. increased linearly with ESd-1. C. ΔES was encoded in dmPFC (MNI = [-S 50 2c]), vmPFC (right, MNI = [3 44 -2c]; left, MNI = [-c c0 -15]), and ventral striatum (right, MNI = [12 12 -S]; left, MNI = [-12 4 -14]). Decreased ΔES was associated with activation in preSMA (MNI = [-27 2 54]). D. dmPFC and bilateral striatal activity increased linearly with ΔES. E. A whole-brain PPI analysis used the ESd-1 time series in aPFC (MNI = [0 c1 15]) as a seed interacting with current evidence strength (ESd). Increasing discrepancy between ESd and ESd-1, synonymous with increasing prediction error, was associated with greater aPFC connectivity to dorsal ACC (right, MNI = [0 c0 10]; left, MNI = [-4 45 3]), overlapping with ΔES-related activation. All activations are cluster-extent FWE-corrected at p < .001. Normalised mean beta values are shown on the y-axes.

Next, we found evidence-strength prediction error (ΔES) signals in a network resembling canonical reward prediction-error circuitry ^30,31^. This included medial prefrontal clusters encompassing ACC (dmPFC, MNI = [-9 50 26], t = 7.70, pFWE < .05; Fig. 3C) and bilateral striatal activation, including ventral striatum (right: MNI = [12 12 -9], t = 5.49, SVC pFWE < .001**; left: MNI = [-12 4 -14], t = 7.31, SVC pFWE < .001**; Fig. 3C). We found negative prediction errors in posterior dmPFC/preSMA (MNI = [-27 2 54], t = 7.33, FWE p < .001; Fig. 3C), consistent with previous observations of inverse reward prediction errors in this region ^32–34^. Thus, our findings suggest that evidence-strength prediction errors share both computational form and neural architecture with traditional reward prediction errors. Both left dmPFC and bilateral NAcc activity scaled linearly with changes in evidence strength (dmPFC: b = 0.07, 95% CI [0.04, 0.09], p < .001; NAcc: b = 0.09, 95% CI [0.04, 0.15], p = .001; Fig. 3D.

### Functional connectivity supports computation of evidence-strength prediction errors

One key expectation for a prediction error is that it integrates information from both prior expectations (in our case, ESd-1) and the current information. To test whether ΔES indeed links these constituents, we conducted a whole-brain psychophysiological interaction (PPI) analysis, using peak aPFC ESd-1 activity as the physiological variable and the currently observed evidence (ESd) as the psychological variable. This ROI was selected as the corresponding region was uniquely associated with ESd-1.

We found a negative interaction, analogous to a prediction-error signal, with increased connectivity in medial prefrontal regions encompassing bilateral ACC (right: MNI = [0 60 10], t = 5.76, FWE p < .001; left: MNI = [-4 45 3], t = 3.35, FWE p < .001; Fig. 3E). These regions closely mirrored the main ΔES contrast, providing convergent evidence that evidence-strength prediction errors arise from interactions between neural representations of prior evidence and currently observed information.

### Urgency recruits dmPFC and locus coeruleus

Investigating urgency, the signal that increases the tendency to commit as time elapses, independent of the evidence, we found a network markedly distinct from the evidence-related networks (see Supplementary Fig. S4). Urgency was associated with increased right dmPFC (MNI = [8 38 22], t = 11.23, pFWE (peak) < .05) and LC activity (right, MNI = [6-32 -18], t = 6.55, pFWE < .001; left, MNI = [-4 -37 -22], t = 6.80, pFWE < .001; confirmed via SVC using an anatomical LC mask, cluster-level FWE-corrected p < .05; see Methods; Fig. 4A). Trial-by-trial BOLD analyses confirmed linear increases in dmPFC activity with trial number (b = 0.24, 95% CI [0.17, 0.32], p < .001) and a trend-level increase in bilateral LC (b = 0.08, 95% CI [-0.005, 0.16], p = .06). In our standard definition of urgency, we characterised the signal as linearly increasing from the beginning of each game (see Methods). However, if this signal should be akin to collapsing boundaries known from sequential sampling models ^21,35^, one would expect that this signal would increase particularly when approaching the termination stage. We thus re-analysed the data by characterising it as a distance to the (theoretical) termination, which incorporates the horizon condition too. We indeed find similar regions to our main analysis (see Supplementary Fig. S3 and Table S3). This indeed supports a notion that also in information gathering an urgency forms a strong pull towards making a decision, capitalising on a network which is akin to the noradrenaline system, in line with recent pharmacological evidence ^16^.

**Figure 4:**
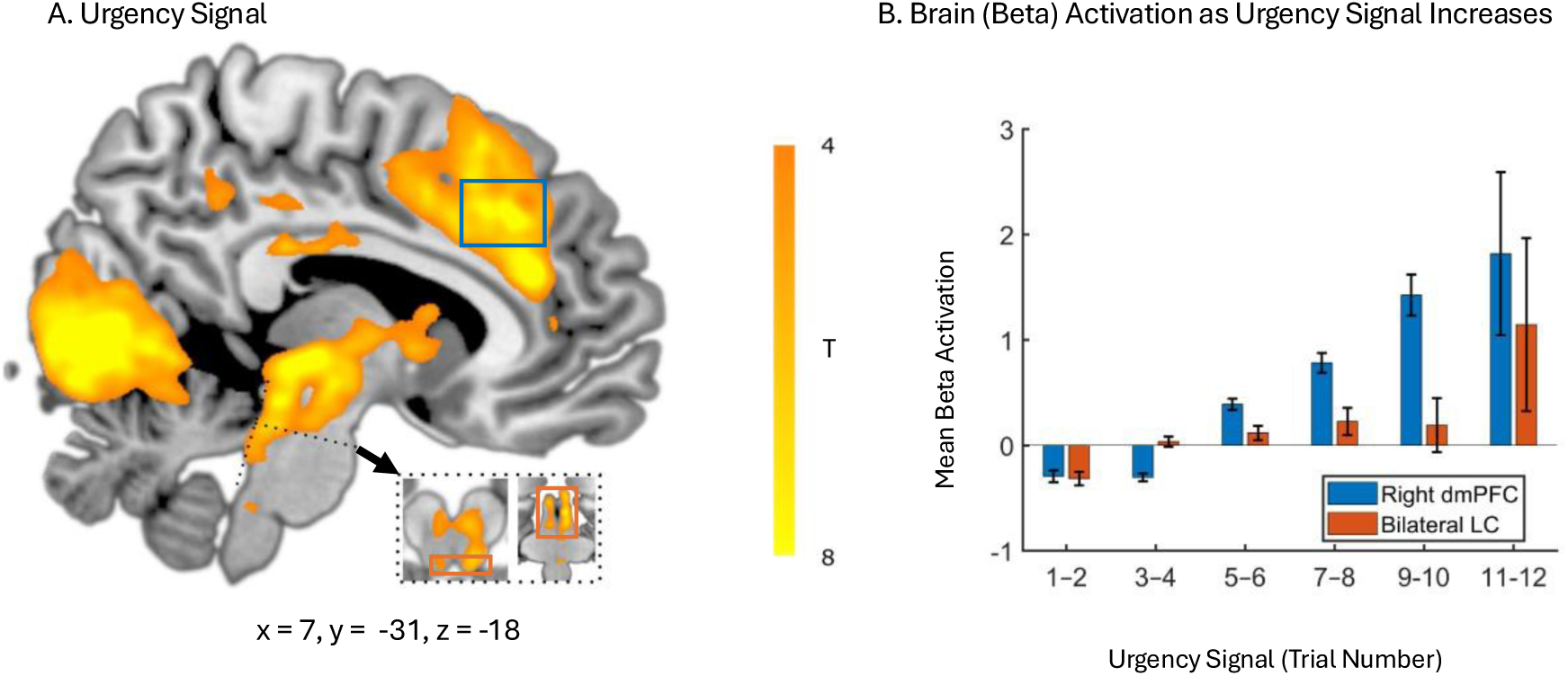
Neural mechanisms subserving the urgency signal. A. Urgency was encoded in right dmPFC (MNI = [8 38 22]) and bilateral LC (right, MNI = [6 -32 -18]; left, MNI = [-4 -37 -22]). All reported activations showed cluster-extent FWE-corrected p < .001. B. Activity in right dmPFC and LC increased with urgency. Normalised mean beta activity values are presented on the y-axis.

## Discussion

Information gathering is a critical, yet understudied, component of decision making, often found to be impaired in mental health conditions. Here, we identify several distinct cognitive constituents that govern information gathering, which recruit different frontal networks, all culminating in a common decision cessation signal in dmPFC.

One key behavioural insight gained from the present study is that gathered information is not weighted equitably, which would, otherwise, constitute an optimal solution. Rather, we find a strong recency bias where most recent information is systematically over-weighted. This extends prior work indicating similar biases in perceptual decision making ^17,18,36,37^. Critically, we found this was driven by a prediction error-like signal (ΔES), signalling the extent to which the most recent information is surprising. Our neuroimaging findings further strengthen this proposed mechanism. Our results reveal distinct, yet interlinked, networks for ΔES and previously gathered information (ESd-1). This suggests that prediction error-like signals are processed separately and used to update cumulative evidence signals in medial prefrontal regions, with such cumulative evidence building the foundation for computing the surprise signal.

Furthermore, we observed that ΔES elicited a network akin to classic reward prediction errors, encompassing ventral striatum and dmPFC. Whilst dmPFC also responded to other cognitive constituents in partially overlapping regions, we found the ventral striatum exclusively tracked ΔES. It is critical to note that unlike classical reward prediction error signals in ventral striatum ^38–42^, in our paradigm there were no rewards delivered at the time of ΔES, clearly setting it apart from standard reward prediction error formulations. Our findings are thus aligned with a more general notion of prediction error processing in this network, going beyond processing tied to rewards ^43,44^ .

We found that the accumulated (prior) information was encoded in the medial prefrontal wall encompassing aPFC and vmPFC. This is significant as complementary research on value-based decision making ^28,45–47^, as well as on latent cognitive state representations ^48–51^ suggests these regions reflect subjective, latent states, either in cognitive or value-based reference spaces. It is striking, therefore, that the computation of the accumulated evidence space also activates similar structures. This also aligns with rodent work showing accumulated sampling information in perceptual decision making in the infralimbic cortex ^52^. Ultimately, we found that this region integrated the accumulated evidence up to the current sample (ESd) with the prior expectation (ESd-1) to compute a prediction error-like signal in the same dmPFC region where we identified ΔES, providing further evidence that the surprise signal builds on a combination of prior expectations and newly accumulated evidence. Distinct from these information-related signals, we found an information-agnostic urgency signal to be computed in a circuit encompassing LC and dmPFC. These regions are well known to constitute a core noradrenergic network with LC hosting most noradrenergic neurons in the brain and dmPFC being a primary target of these projections ^53–55^. The potential contribution of noradrenaline is further supported by our recent evidence showing that noraderenegic agents modulate information gathering ^16,56,57^. Interestingly, this somewhat departs from the neural circuits suggested for computing urgency and decision thresholds in perceptual decision making, which has primarily implicated fronto-striatal loops including basal ganglia and premotor cortex ^58–60^ or subthalamic nuclei ^27,61^. It is quite possible that, albeit similar in its cognitive function, an urgency signal might be recruiting different neural circuits depending on the formulations, as the timescales are markedly different between perceptual decision making and information gathering. Interestingly, the notion of noradrenaline encoding urgency aligns well with dominant theories of LC where its proposed role is in signalling task utility ^62^, whereby reduced utility promotes disengagement from an ongoing task process. It further aligns with evidence for dmPFC activity scaling with advantageous action timing in competitive decision-making tasks ^63^, suggesting that this region supports strategic time-based information gathering.

Finally, we identified the dmPFC as an integrative hub for information gathering, holding partially overlapping information about the specific cognitive constituents, such as evidence and urgency, as well as a cumulative signal integrating across neurocognitive constituents of information gathering. This finding accords with evidence that dmPFC activity reflects the value of incoming information relative to what is already known ^12^, suggesting continuous monitoring and updating of the evidence landscape to guide sampling cessation. Our findings extend this account by showing that dmPFC not only responds to incoming evidence but also tracks accumulating pressure to commit, integrating information content and urgency.

Our findings demonstrate that a seemingly simple behavioural process, such as information gathering, results from a complex interplay between distinct cognitive contributors, each recruiting distinct neural circuits that converge into an integrated signal determining the extent of information gathering. Our findings demonstrate that these cognitive constituents build on processes (such as prediction errors) and networks that have been identified in different contexts, such as reward processing, suggesting they serve more general-purpose functions. Our findings further yield insight into potential pathological mechanisms that underlie stark information-gathering biases in mental health conditions, such as jumping-to-conclusions in schizophrenia ^64–66^ and indecisiveness in obsessive-compulsive disorder (OCD);^67–70^. Tracking the exact neurocognitive mechanisms that go awry in these conditions could allow us to build novel, targeted interventions.

## Methods

### Participants

Twenty-nine healthy volunteers were recruited from a local volunteer pool and compensated hourly, with a performance-dependent bonus. One participant was excluded due to excessive movement during scanning. The study was approved by the University of Tübingen Ethics Committee (268/2023BO2) and participants provided written informed consent.

### Information gathering task

Participants performed an fMRI information-gathering task building on a task we developed previously (see del Río et al. (2025) for further details). On each game, participants chose which of two stimuli was more abundant overall. Stimuli were blue zircons (blue squares) or gold nuggets (yellow squares) (Fig. 1A).

Each game comprised 4-12 consecutive draws. Before the first draw, a grey square was displayed for 1-4 s (mean = 2 s; Poisson-distributed). Draw number was pseudo-randomised across two conditions: long horizon (8-12 draws) and short horizon (4-8 draws) (Fig. 1B), which were indicated by frame colour (pink or green) and learned prior to the fMRI.

Each draw presented five stimuli for 2 s, followed by an inter-draw interval of 1-6 s (mean = 2 s). The horizon cue remained visible throughout each game, allowing participants to estimate expected game length without revealing the exact number of draws.

Participants had until the final draw to indicate which stimulus they expected to be most abundant across all draws in that game.

Correct responses earned 2 points, and incorrect responses lost 2 points. If no response was made before game termination, participants lost 1 point. Feedback was displayed for 1 s before the next game. A progress bar indexed accumulated reward and reset after reaching the bonus threshold.

### Behavioural analysis

To characterise information-gathering constituents, we used a mixed-effects logistic regression predicting whether participants stopped sampling or gathered additional information on each trial. Predictors were horizon, urgency (number of draws, Nd), total evidence strength from the previous draw (ESd-1), and evidence-strength prediction error (ΔES), building on our previous work (del Río et al., 2025; Fig. 1C). The model-predicted likelihood for each sample was defined as the decision-commitment signal (DCS). Horizon was coded as a binary variable distinguishing short from long horizons, with long horizons providing larger sampling windows.

Urgency was defined as the number of draws observed so far. Evidence strength (ES) was quantified separately for each stimulus. For stimulus A, cumulative ES at draw d was the sum of all A instances observed up to that draw.

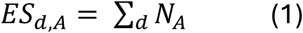

Similarly for stimulus B:

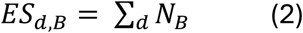

Total evidence strength (ES) was defined as a single measure indexing the extent to which accumulated evidence favoured one option. ESd,majority denoted cumulative evidence for the stimulus predominant at draw d, and ESd,minority denoted cumulative evidence for the alternative stimulus. At each draw, ESd-1 was defined as their lagged difference.

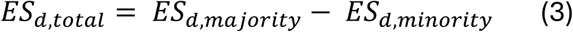

ΔES was defined as the change in ES from the previous draw to the current draw.

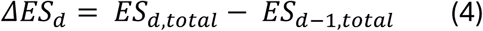

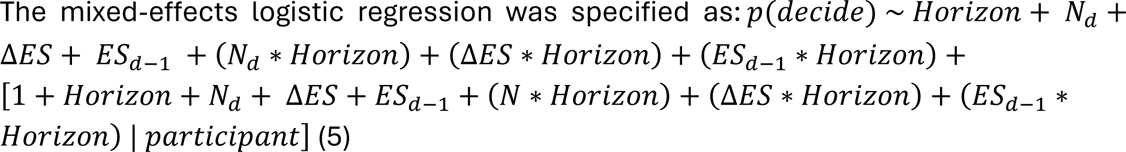

### fMRI data acquisition and preprocessing

Participants performed the task during functional MRI on a 3T Siemens Prisma scanner with a 64-channel head coil at the University Hospital Tübingen. High-resolution anatomical images were acquired using a T1-weighted MPRAGE sequence (TR = 2530 ms, TE = 3.34 ms, voxel size = 0.8 mm3, 176 sagittal slices). Functional images were acquired using a multiband T2*-weighted EPI sequence (TR = 1500 ms, TE = 20 ms, flip angle = 65°, voxel size = 2.4 mm3, 60 transversal slices) following ^71^ scanning procedure. Fieldmaps were acquired with a gradient-echo sequence (TR = 1020 ms, TE1 = 10 ms, TE2 = 12.45 ms, voxel size = 3.0 x 3.0 x 2.0 mm3) to correct EPI field-strength inhomogeneity.

Physiological signals were recorded during scanning (respiration belt and pulse oximeter; BIOPAC Systems, Inc.). Thirteen regressors were derived using RETROICOR ^72^: six cardiac regressors, six respiratory regressors, and one cardiorespiratory interaction regressor. Regressors were estimated with the PhysIO toolbox ^73^ and included for physiological noise correction in the following fMRI analysis.

Functional and structural MRI analyses were performed using SPM12. The first six scans of each session were discarded. Voxel displacement maps were calculated for each EPI run using the SPM12 FieldMap toolbox and applied during realignment and unwarping, correcting each EPI volume for head motion and geometric distortion.

The mean functional image was coregistered to each participant’s T1-weighted image, and this alignment was applied to all functional images. T1-weighted images were segmented into tissue probability maps, and resulting deformation fields normalised functional data to MNI space. Normalised EPI images were smoothed using a 6-mm FWHM isotropic Gaussian kernel.

### fMRI data analysis

fMRI analyses identified neural correlates of behavioural processes underlying termination of information sampling and subsequent choice commitment.

Analyses proceeded hierarchically. We first modelled predicted choice probability to identify regions tracking the overall decision signal and then decomposed this signal into constituent behavioural components to determine their specific neural correlates.

In the first model, predicted choice probability on each trial was entered as a parametric regressor in the GLM. Cue-related activity was controlled by including cue onset as the base regressor for the parametric modulator, with orthogonalisation deactivated to allow each regressor to explain unique variance. Missing values for ΔEV, Esd-1, and p(decide) arose from the absence of prior evidence on the first draw. For p(decide), these were replaced with the minimum z-scored value, reflecting the reduced likelihood of committing to a decision at this stage. For ΔEV and Esd-1, the minimum was not an appropriate substitute, as it carries a distinct meaning for these variables; missing values were instead replaced with the corresponding variable mean (Fig. 2A).

Subsequent contrasts disentangled feature-specific contributions to decision making. Trial-wise onsets and parametric modulators were derived from behavioural data. Z-scored ΔES, urgency, and ESd were included as parametric modulators of cue onsets, with orthogonalization deactivated. Cue regressors controlled for visual task-related activity (Figs. 3A, C and Fig. 4A).

Additional analyses included response and horizon regressors, as described in the Supplementary Figs S1-S3.

Each task block was modelled as a separate run to account for baseline signal differences. First-level contrast images were entered into second-level random-effects analyses and submitted to second-level t-tests across participants. Positive and negative effects were examined separately. Statistical significance was assessed using a voxel-wise threshold of p < .001 (uncorrected) and cluster-level FWE correction at p < .05 (Figs. 2 and 3); peak voxels within significant clusters are reported with FWE-corrected peak-level p-values.

### Region-of-interest analysis

Two regions of interest were defined a priori for small-volume correction (SVC) of the LC and ventral striatal/NAcc effects. The LC was defined as an a priori ROI using the consensus mask from Mäki-Marttunen and Espeseth (2020), resliced to functional resolution with the mean BOLD image as reference and thresholded at p > .5 to retain high-confidence LC voxels. For ventral striatum/nucleus accumbens (NAcc), whole-brain analysis revealed a large striatal cluster peaking in dorsal striatum. Given our a priori hypothesis concerning ventral striatal prediction-error signalling and cluster extension into NAcc, SVC used the probabilistic striatal atlas from Pauli et al. (2018), thresholder at p > .5. For both ROIs, peak-level FWE-corrected p values are reported within the restricted search volume.

### Trial-by-trial ROI analysis

For illustrative analyses, functionally defined ROIs were centred on peak activations from relevant contrasts (Table 1) with a 3-mm spherical radius. Trial-by-trial activations were extracted by fitting a GLM containing cue events only, yielding beta maps for each trial. ROI-overlapping beta values were then extracted for further analysis (Fig. 2A, Figs 3B, D and Fig.4B).

### Psychophysiological interaction analysis

To assess whether ΔES regions were functionally coupled with regions representing prior evidence, we conducted a PPI analysis (Friston et al., 1997). The seed was a 6-mm spherical ROI in anterior medial prefrontal cortex (aPFC) at MNI = [0 61 15], which correspond to the aPFC, a region which was solely linked to ESd-1. The seed time series was extracted for each participant and entered into a PPI model interacting with trial-wise current evidence strength (ESd), normalised across trials.

In the first-level GLM, ESd also served as a cue-onset parametric modulator. The resulting interaction term was included as a regressor in a whole-brain GLM. We examined the negative interaction term to identify connectivity increases when current evidence (ESd) exceeded accumulated prior evidence (ESd-1).

## Acknowledgments

We thank the Core Facility MRI of the Faculty of Medicine Tübingen for providing excellent technical support in installing and troubleshooting MR sequences. We also thank Luca Kosina for her work on the task implementation. TUH has received funding from the Wellcome Trust (316955/Z/24/Z), the European Research Council (ERC) under the European Union’s Horizon 2020 research and innovation programme (grant agreement No 946055), and the Carl-Zeiss-Stiftung. This work was supported by the Alexander von Humboldt foundation, (more) precisely the Alexander-von-Humboldt-Professorship award to Peter Dayan. Funding by the Deutsche Forschungsgemeinschaft (DFG - German Research Foundation, project number 460704019) is gratefully acknowledged. TUH consults for limbic ltd and holds share options in the company, which is unrelated to the current project.

## Notes

### Competing Interest Statement

The authors have declared no competing interest.

