## Supplementary Material for "Dorsomedial prefrontal cortex acts as an integrative hub during information gathering"

Content:


- Figure S1: Whole-brain activations for decision-related regressors controlled for motor response
- Figure S2: Neural correlates of decision-related regressors controlling for horizon length
- Figure S3: Whole-brain activations for decision-related regressors replacing urgency by termination proximity (horizon-corrected)
- Figure S4: Distinct neural networks implicated in urgency and evidence accumulation
- Table S1: Peak activations for decision-related regressors controlling for motor response
  Table S2: Peak activations for decision-related regressors controlling for horizon length
  Table S3: Peak activations for decision-related regressors using termination proximity

**Figure S1: Whole-brain activations for decision-related regressors controlled for motor response.** A. Decision-commitment was still encoded (yellow) in right dmPFC (MNI = [4 28 39]) and right LC (MNI = [4 -36 -21]), both showing cluster-extent FWE-corrected p < .001. Lowered p(decide) (blue blobs) was associated with activation in left vmPFC (MNI = [-6 44 -14]). B. ΔES was encoded in aPFC (right, MNI = [6 56 9]; left, MNI = [-3 56 -22]), vmPFC (right, MNI = [2 34 -15]; left, MNI = [-6 48 -15]), and ventral striatum (right, MNI = [12 12 -9]; left, MNI = [-10 4 -14]), confirmed via SVC using an anatomical NAcc mask, cluster-level FWE-corrected p < .05. C. The urgency signal was encoded in right dmPFC (MNI = [8 38 22], pFWE(peak) < .05) and LC (right, MNI = [4 -34 -20], t = 4.62; left, MNI = [-4 -36 -21], t = 5.23), confirmed via SVC using an anatomical LC mask, cluster-level FWE-corrected p < .05 (see Methods). D. ESd-1 was primarily encoded in bilateral dmPFC (right, MNI = [15 60 34]; left, MNI = [-10 51 40]), cluster-level FWE-corrected p < .05.

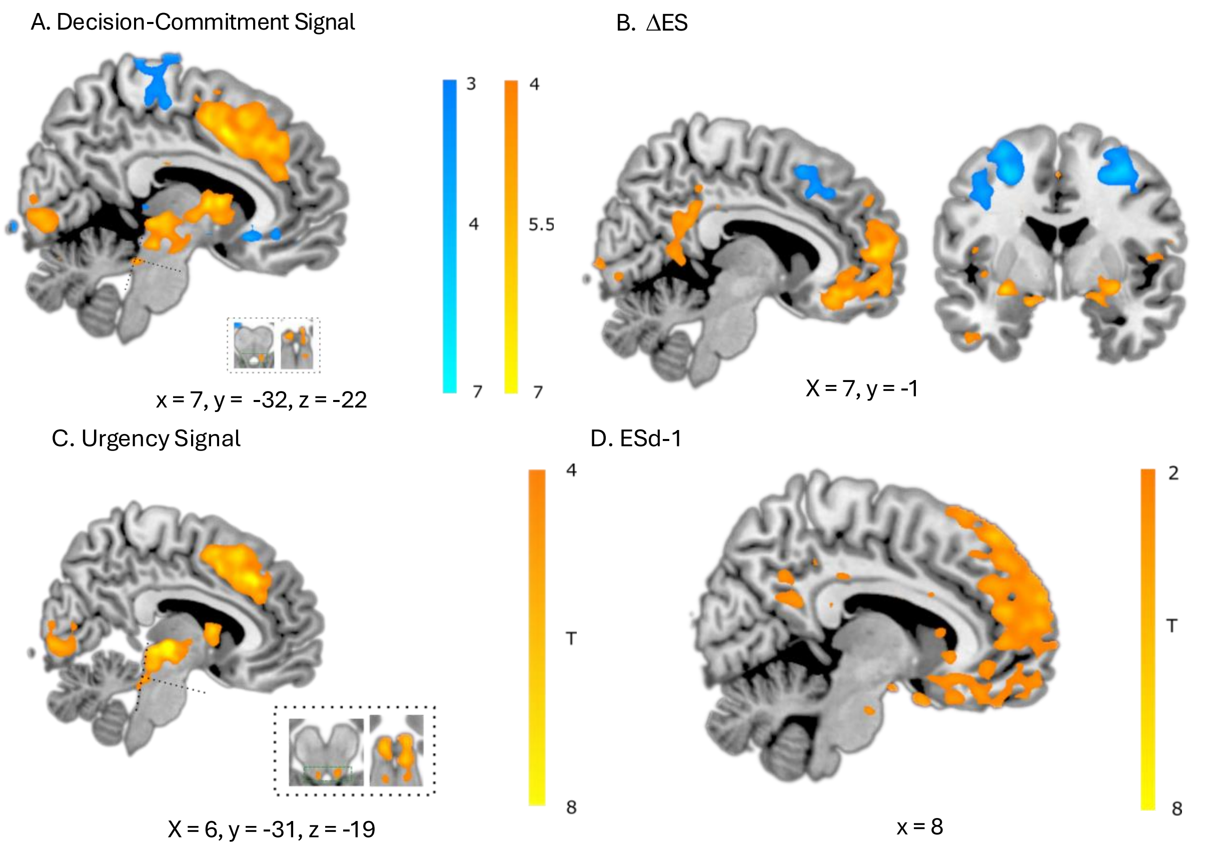

**Figure S2: Neural correlates of decision-related regressors controlling for horizon length.** A. ESd-1 was primarily encoded in bilateral dmPFC (right, MNI = [9 42 52]; left, MNI = [-14 48 42]), with cluster-level FWE correction ($p < .05$). B. ΔES was encoded in bilateral dmPFC (right, MNI = [10 42 52]; left, MNI = [-15 46 38]) and bilateral NAcc (right, MNI = [12 12 -9]; left, MNI = [-12 4 -12]), confirmed via SVC using an anatomical NAcc mask, cluster-level FWE-corrected $p < .05$ (see Methods). Blue blobs indicate deactivations. C. The urgency signal was encoded in right dmPFC (MNI = [6 38 22], pFWE(peak) < .05) and bilateral LC (right, MNI = [4 -34 -20], $t = 4.78$; left, MNI = [-4 -36 -21], $t = 5.03$), confirmed via SVC using an anatomical LC mask, cluster-level FWE-corrected $p < .05$ (see Methods).

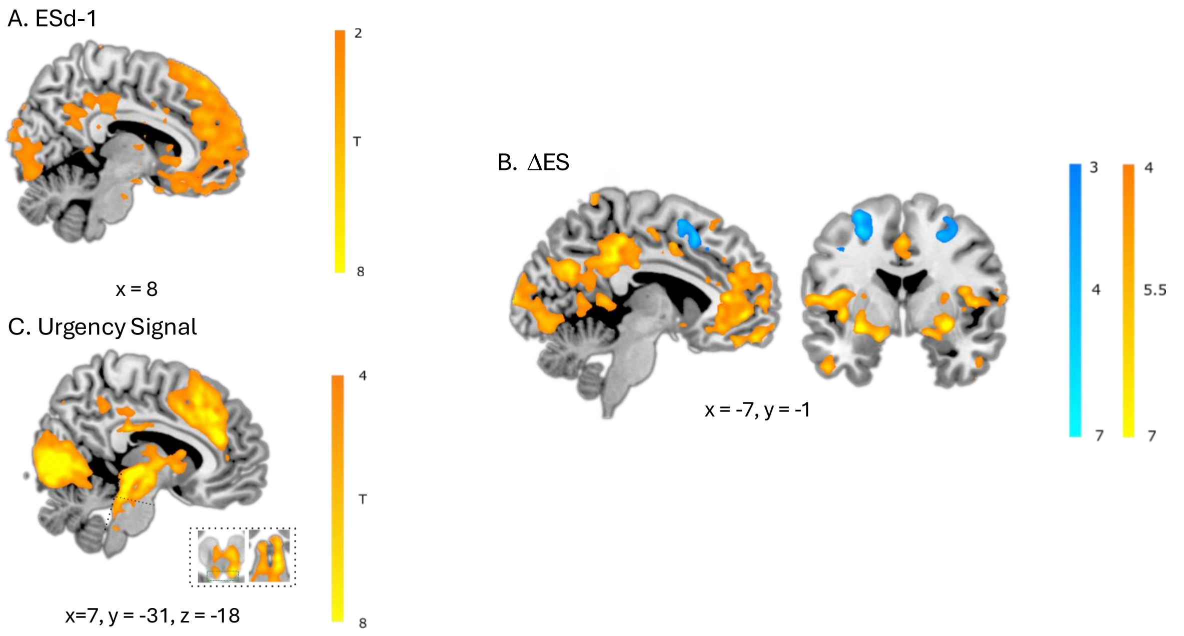

**Figure S3: Whole-brain activations for decision-related regressors replacing urgency by termination proximity (horizon-corrected).** A. ESd-1 was primarily encoded in bilateral dmPFC (right, MNI = [9 42 52], t = 6.77; left, MNI = [-14 48 42], t = 6.27; ‡p < .0001 uncorrected) and vmPFC (right, MNI = [2 36 -20], t = 5.15, NS). B. ΔES was encoded in bilateral dmPFC (right, MNI = [10 42 52], t = 9.24; left, MNI = [-15 46 38], t = 8.03) and bilateral NAcc (right, MNI = [12 12 -9], t = 5.28; left, MNI = [-12 4 -12], t = 7.27), confirmed via SVC using an anatomical NAcc mask, cluster-level FWE-corrected p < .05 (see Methods). Blue blobs indicate deactivations. C. Termination proximity was encoded in right dmPFC (MNI = [6 38 22], t = 10.97, pFWE(peak) < .05) and bilateral LC (right, MNI = [4 -34 -20], t = 4.51; left, MNI = [-4 -36 -26], t = 4.73), confirmed via SVC using an anatomical LC mask, cluster-level FWE-corrected p < .05 (see Methods).

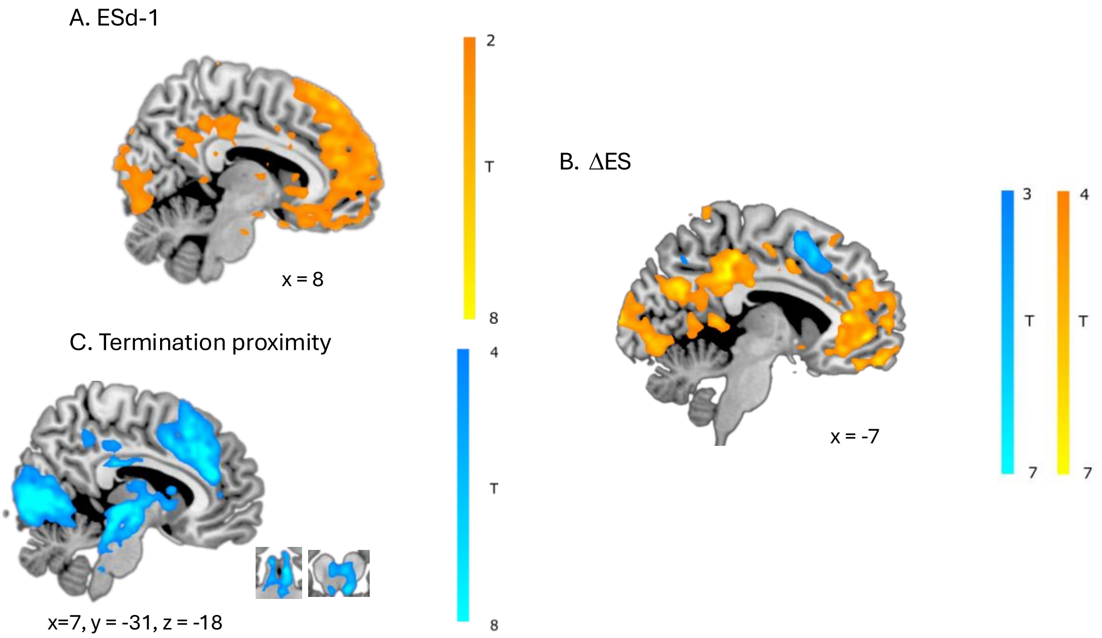

**Figure S4: Distinct neural networks implicated in urgency and evidence accumulation.** Evidence-related networks (ΔES, green; ESd-1, red) are distinct from the urgency network (blue). All three networks overlap in the dmPFC.

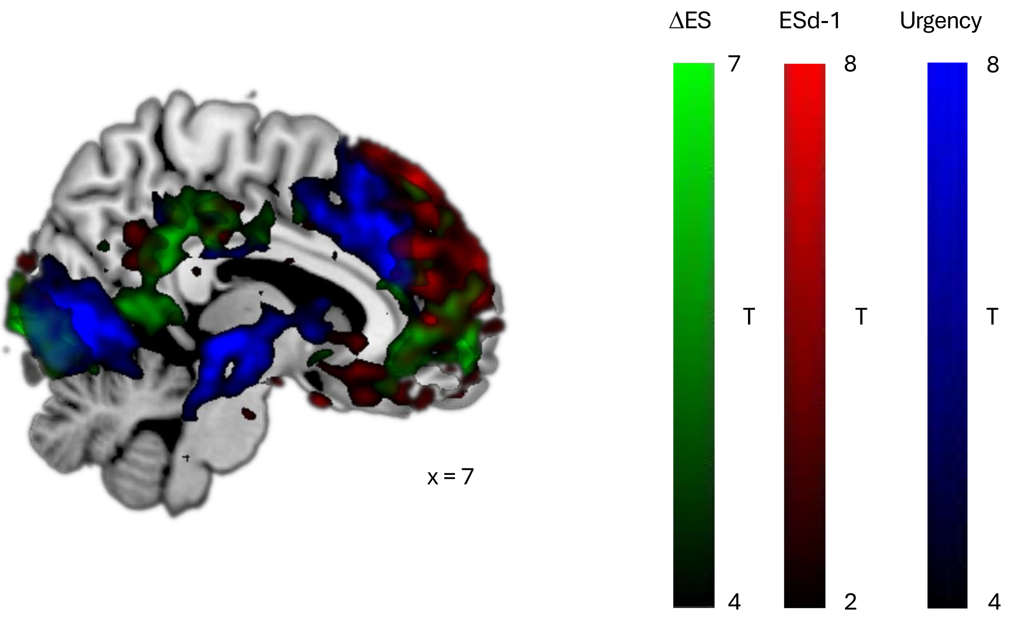

**Table S1: Peak activations for decision-related regressors controlling for motor response.** Clusters were defined using a voxel-wise threshold of $p < .001$ (uncorrected), with cluster-level FWE correction ($p < .05$), unless otherwise indicated.

* pFWE (peak) < .05; ** pFWE (peak) < .05, SVC (LC: Mäki-Marttunen & Espeseth, 2020; NAcc: Pauli et al., 2018); ‡ p < .0001 (uncorrected), NS Non-significant

Abbreviations: dmPFC = dorsomedial prefrontal cortex; vmPFC = ventromedial prefrontal cortex; LC = locus coeruleus; aPFC = anterior prefrontal cortex.

| Contrast | Region | Hemisphere | Cluster Size (voxels) | X | Y | Z | T Score |
|---|---|---|---|---|---|---|---|
| **DCS** | dmPFC | Right* | 45 | 4 | 28 | 39 | 7.68 |
| | LC | Right** | 4 | 4 | -36 | -21 | 3.90 |
| **-DCS** | vmPFC | Right NS | 82 | 12 | 42 | -8 | 4.10 |
| | | Left | 866 | -6 | 44 | -14 | 5.01 |
| **$ES_{d-1}$** | dmPFC | Right | 5194 | 15 | 60 | 34 | 5.42 |
| | | Left | | -10 | 51 | 40 | 6.03 |
| **ΔES** | aPFC | Right‡ | 7324 | 6 | 56 | 9 | 7.97 |
| | | Left‡ | | -3 | 56 | -22 | 7.41 |
| | vmPFC | Right‡ | 7324 | 2 | 34 | -15 | 8.49 |
| | | Left‡ | | -6 | 48 | -15 | 7.41 |
| | Nacc** | Right | 6 | 12 | 12 | -9 | 3.95 |
| | | | 9 | 4 | 8 | -6 | 3.81 |
| | | Left | 40 | -10 | 4 | -14 | 5.10 |
| **Urgency** | dmPFC | Right* | 107 | 8 | 38 | 22 | 11.23 |
| | LC | Right** | 2 | 4 | -34 | -20 | 4.62 |
| | | | 4 | 6 | -36 | -26 | 4.24 |
| | | Left** | 7 | -4 | -36 | -21 | 5.23 |

**Table S2: Peak activations for decision-related regressors controlling for horizon length.** Clusters were defined using a voxel-wise threshold of $p < .001$ (uncorrected), with cluster-level FWE correction ($p < .05$), unless otherwise indicated.

* pFWE (peak) < .05; ** pFWE (peak) < .05, SVC (LC: Mäki-Marttunen & Espeseth, 2020; NAcc: Pauli et al., 2018); ‡ p < .0001 (uncorrected), NS Non-significant

Abbreviations: dmPFC = dorsomedial prefrontal cortex; vmPFC = ventromedial prefrontal cortex; LC = locus coeruleus; aPFC = anterior prefrontal cortex.

| Contrast | Region | | Cluster Size (voxels) | X | Y | Z | T Score |
|---|---|---|---|---|---|---|---|
| **$ES_{d-1}$ with horizon** | dmPFC‡ | Right | 3782 | 9 | 42 | 52 | 6.77 |
| | | Left | | -14 | 48 | 42 | 6.27 |
| **ΔES with horizon** | dmPFC* | Right | 91 | 10 | 42 | 38 | 9.24 |
| | | Left | 13 | -15 | 46 | 38 | 8.03 |
| | Nacc** | Right | 94 | 12 | 12 | -9 | 5.28 |
| | | Left | 128 | -12 | 4 | -12 | 7.27 |
| **Urgency** | dmPFC | Right* | 937 | 6 | 38 | 22 | 10.97 |
| | LC | Right** | 3 | 4 | -34 | -20 | 4.78 |
| | | | 2 | 6 | -38 | -26 | 4.23 |
| | | | 1 | 4 | -36 | -22 | 3.56 |
| | | Left** | 7 | -4 | -36 | -21 | 5.03 |

**Table S3: Peak activations for decision-related regressors using termination proximity.** Clusters were defined using a voxel-wise threshold of $p < .001$ (uncorrected), with cluster-level FWE correction ($p < .05$), unless otherwise indicated.

* pFWE (peak) < .05; ** pFWE (peak) < .05, SVC (LC: Mäki-Marttunen & Espeseth, 2020; NAcc: Pauli et al., 2018); ‡ p < .0001 (uncorrected), NS Non-significant

Abbreviations: dmPFC = dorsomedial prefrontal cortex; vmPFC = ventromedial prefrontal cortex; LC = locus coeruleus; aPFC = anterior prefrontal cortex.

| Contrast | Region | | Cluster Size (voxels) | X | Y | Z | T Score |
|---|---|---|---|---|---|---|---|
| **$ES_{d-1}$** | dmPFC‡ | Right | 3782 | 9 | 42 | 52 | 6.77 |
| | | Left | | -14 | 48 | 42 | 6.27 |
| | vmPFC‡ | Right NS | 148 | 2 | 36 | -20 | 5.15 |
| **ΔES** | dmPFC* | Right | 91 | 10 | 42 | 52 | 9.24 |
| | | Left | 13 | -15 | 46 | 38 | 8.03 |
| | Nacc** | Right | 94 | 12 | 12 | -9 | 5.28 |
| | | Left | 128 | -12 | 4 | -12 | 7.27 |
| **-Termination proximity** | dmPFC | Right* | 937 | 6 | 38 | 22 | 10.97 |
| | LC** | Right | 2 | 4 | -34 | -20 | 4.51 |
| | | | 2 | 6 | -36 | -26 | 3.99 |
| | | Left | 6 | -4 | -36 | -26 | 4.73 |
